# Structural characterization of the Sel1-like repeat protein LceB from *Legionella pneumophila*

**DOI:** 10.1101/2023.07.03.547437

**Authors:** Tiffany V. Penner, Neil Lorente Cobo, Deepak T. Patel, Dhruvin H. Patel, Alexei Savchenko, Ann Karen C. Brassinga, Gerd Prehna

## Abstract

*Legionella* are freshwater Gram-negative bacteria that in their normal environment infect protozoa. However, this adaptation also allows *Legionella* to infect human alveolar macrophages and cause pneumonia. Central to *Legionella* pathogenesis are more than 330 secreted effectors, of which there are 9 core effectors that are conserved in all pathogenic species. Despite their importance, the biochemical function of several core effectors remains unclear. To address this, we have taken a structural approach to characterize the core effector of unknown function LceB, or Lpg1356, from *Legionella pneumophila.* Here we solve an X-ray crystal structure of LceB using an AlphaFold model for molecular replacement. The experimental structure shows that LceB adopts a Sel1-like repeat fold as predicted. However, the crystal structure captured multiple conformations of LceB all of which differed from the AlphaFold model. Comparison of the predicted model and the experimental models suggests that LceB is highly flexible in solution. Additionally, molecular analysis of LceB using its close structural homologues reveals sequence and structural motifs of known biochemical function. Specifically, LceB harbors a repeated KAAEQG motif that both stabilizes the Sel1-like repeat fold and is known to participate in protein-protein interactions with eukaryotic host proteins. We also observe that LceB forms several higher-order oligomers in solution. Overall, our results have revealed that LceB has conformational flexibility, self-associates, and contains a molecular surface for binding a target host-cell protein. Additionally, our data provides structural insights into the Sel1-like repeat family of proteins that remain poorly studied.

## INTRODUCTION

*Legionella* species are Gram-negative bacteria ubiquitous in freshwater environments^1^ where they parasitize protozoa^2^. By evolving with protozoa, *Legionella* has established survival and replication mechanisms to persist within eukaryotic cells such as amoeba. Specifically, the similarity of an amoeba to macrophages has primed *Legionella* to be a serious pathogen capable of infecting human alveolar macrophages^2, 3^. Infection of human alveolar macrophages can occur by inhalation of contaminated water aerosols. This may result in Legionnaires’ disease which is characterized by severe pneumonia primarily in elderly and/or immunocompromised individuals^4–8^,. Prompted by phagocytosis, Legionellae utilize the Dot/Icm Type IV secretion system (T4SS) to translocate effector proteins into the host cytosol to establish a replicative niche known as the *Legionella*-containing vacuole (LCV)^9, 10^. Vesicles from the smooth endoplasmic reticulum (ER) fuse with the LCV membrane and eventually become studded with ribosomes and mitochondria^11^. The LCV evades fusion with lysosomes and maintains a higher pH than vacuoles with formalin-killed *Legionella pneumophila*^12, 13^. To establish and maintain the LCV, T4SS effector proteins are secreted into the host macrophage which are crucial for manipulating host cell signaling pathways and virulence^14, 15^. In *L. pneumophila* alone, more than 330 effectors are known to be translocated^5^. Moreover, bioinformatic studies have predicted that there are over 18,000 T4SS effectors from 58 *Legionella* species, with only nine that are conserved in all species^16, 17^. Given the vast array of effectors the majority of these proteins have not been fully characterized due to host specificity and redundancy^5, 18^. Those T4SS proteins that have been studied, display a variety of functions including evasion from the endocytic maturation pathway, interaction with the ER, kinase signaling, epigenetic regulation, mRNA processing and manipulation of the ubiquitin pathway^5^.

As many *Legionella* effectors directly bind protein targets in their hosts, these effectors often contain tetratricopeptide repeats. Tetratricopeptide repeats (TPRs) were first discovered in the eukaryote *Saccharomyces cerevisiae* gene product CDC23 which has roles in the synthesis of RNA and mitosis^19^. These motifs are composed of 34 loosely conserved, alternating large and small sidechain residues. TPRs are usually found in proteins as 3-16 tandem repeats but may be dispersed throughout the protein^19, 20^. In bacterial species, TPR proteins are known to function in several pathways including bio-mineralization of iron oxides in magnetotactic bacteria^21^, assembly of the outer membrane^22^, natural competence^23, 24^ and pathogenesis^23, 25, 26^ including the movement of virulence factors into host cells^27, 28^. Although TPR proteins participate in a wide variety of biological functions, one commonality is their importance in protein-protein interactions^29, 30^. Given this, TPRs are ubiquitous in nature^29^ and found in all 3 domains of life^31^.

A subtype of TPR proteins exist termed Sel1-like repeat (SLR) proteins, which are a structural variation of the canonical TPR^32^. Although the α-helical conformations adopted by TPR and SLR proteins are similar, there are spatial differences between TPR and SLR proteins. This was first shown in the structure of the SLR protein HcpB from *Helicobacter pylori*^33^. As compared to TPRs, SLRs have 4-12 additional residues in the loop between the two α-helices of a single repeat unit, but 2 fewer residues in the repeat connector^34^. Furthermore, the SLR sequence is longer than the TPR as it consists of 36 to 44 amino acid residues^34^. However, both motifs contain the characteristic pattern of large and small residue sidechains, including tyrosine. For example, according to the SMART database^20^ the canonical SLR motif is 3A-7L-8G-11Y-14G-16G-20D-24A-31A-32A-35G, and the canonical TPR motif is 4W-7L-8G-11Y-20A-24F-27A-32P^19^.

SLR proteins were first described in the round-worm *Caenorhabditis elegans*^35^. In this work, the extracellular SEL-1 protein was shown to be a negative regulator of membrane receptor proteins Lin-12 and Glp-1 which are both important for determining the fate of a cell^35^. Although the exact mechanism of SEL-1 is unknown, it is likely that SEL-1 targets the Lin-12 receptor when bound to a ligand for subsequent degradation^36^. Bacterial SLRs have been shown to function in exopolysaccharide synthesis in Pseudomonas^37^, flagellar motility in *Vibrio parahemolyticus*^38^, and infection in *L. pneumophila* by mediating the interaction between the bacteria and its eukaryotic host^27, 39, 40^ Not unlike TPR proteins, SLR proteins have a variety of functions that are part of signal transduction pathways and immunomodulatory functions^34^. However, limited literature exists on SLR proteins as compared to TPR domains due to the lower abundance of the SLR fold. This lack of data includes not only the discovery of biological functions, but experimental SLR protein structures alone and in complex with their biochemical targets^41^.

The *L. pneumophila* genome contains 5 open reading frames that encode SLR proteins^42^, making this pathogen a powerful model to study the biochemistry of the SLR fold. These genes encode the SLR proteins LpnE, EnhC, LidL, Lpg1062, and LceB. The proteins are thought to be secreted virulence factors, however the translocation mechanism of each remains unclear. To the best of our knowledge LpnE is neither translocated by the Type II secretion system (T2SS) or T4SS^42^. There is data that shows LceB is translocated by the T4SS^17^, but LceB also contains a general N-terminal secretory signal (sec-signal) making the overall mechanism of its secretion unclear.

In terms of biological function, each of the 5 *L. pneumophila* SLR proteins has been studied. Both LpnE and EnhC have been extensively characterized and are known to assist in host cellular invasion^39, 40^. Additionally, these two proteins along with LidL are important in the signaling events required for proper trafficking of *L. pneumophilia* within macrophage cells^42, 432^. The protein Lpg1062 is required for growth in *Naegleria gruberi*^3^, is predicted to have 8 SLR folds^32^, and like LceB contains a sec-signal sequence^44^. LceB was recently discovered to be one of the 9 core effectors in *Legionella*^16^. Further investigation showed that a deletion mutant of LceB grew similarly to wildtype strains during competition assays using *Acanthamoeba castellanii* as a host^17^. Furthermore, an elegant study utilizing transposon mutagenesis also concluded that LceB was not important for intracellular growth *in A. castellanii*. This same study also determined that LceB was not required for growth in *Acanthamoeba polyphaga*, *Hartmannella vermiformis*, and *N. gruberi*, but necessary for growth in human U937 macrophages^3^. Thus, although the biochemical function of LceB remains unclear, its requirement for growth in macrophages suggests that it may play a specialized role in *Legionella* pathogenesis of human hosts.

Given that LceB is a secreted SLR protein effector of unknown function, we sought to characterize the structure of LceB to provide insight into its biochemical function at the molecular level. Here we solve an X-ray crystal structure of LceB that reveals an Sel1-like repeat protein in agreement with bioinformatic predictions. Structural analysis of LceB compared to its close homologs reveal several conserved structural features and sequence motifs that provide insight into the role of LceB during *Legionella* infection. Furthermore, LceB crystallized with 3 copies in the asymmetric unit which adopted unique conformations different from the AlphaFold prediction. Overall, our data shows a dynamic SLR *Legionella* effector that contains several structural and sequence motifs also found in functionally characterized bacterial toxins.

## RESULTS AND DISCUSSION

### Overall structure of LceB

To purify LceB for crystallization, residues 1-18 were removed from the expression construct as they are predicted to be a sec-signal^44^. Instead, a construct spanning residues 22-366 from *L. pneumophila* containing an N-terminal 6His-tag was purified and screened for crystallization. Initially, poorly diffracting crystals were obtained. To improve crystallization quality, the buffer for LceB was optimized using nanodifferential scanning fluorimetry (nanoDSF) by monitoring the change in tryptophan fluorescence during thermal denaturation (Fig. S1 and Table S1). This resulted in a significant increase of the LceB T_m_ from ∼63 °C to ∼68 °C by raising the NaCl concentration from 250 mM to 750 mM. LceB was repurified in the optimized buffer, crystallized, and an X-ray crystal structure solved to a resolution of 2.7 Å (Fig. 1 and Table 1). LceB crystallized in space group P3_2_ with three chains in the asymmetric unit (Fig. S2). Analysis of the asymmetric unit by PDBePISA^45^ suggests these are likely crystal packing artifacts as the contact surface for each chain varies with the largest interaction at 644.3 Å^2^ ^45^ (Table S2).

**Figure 1.**
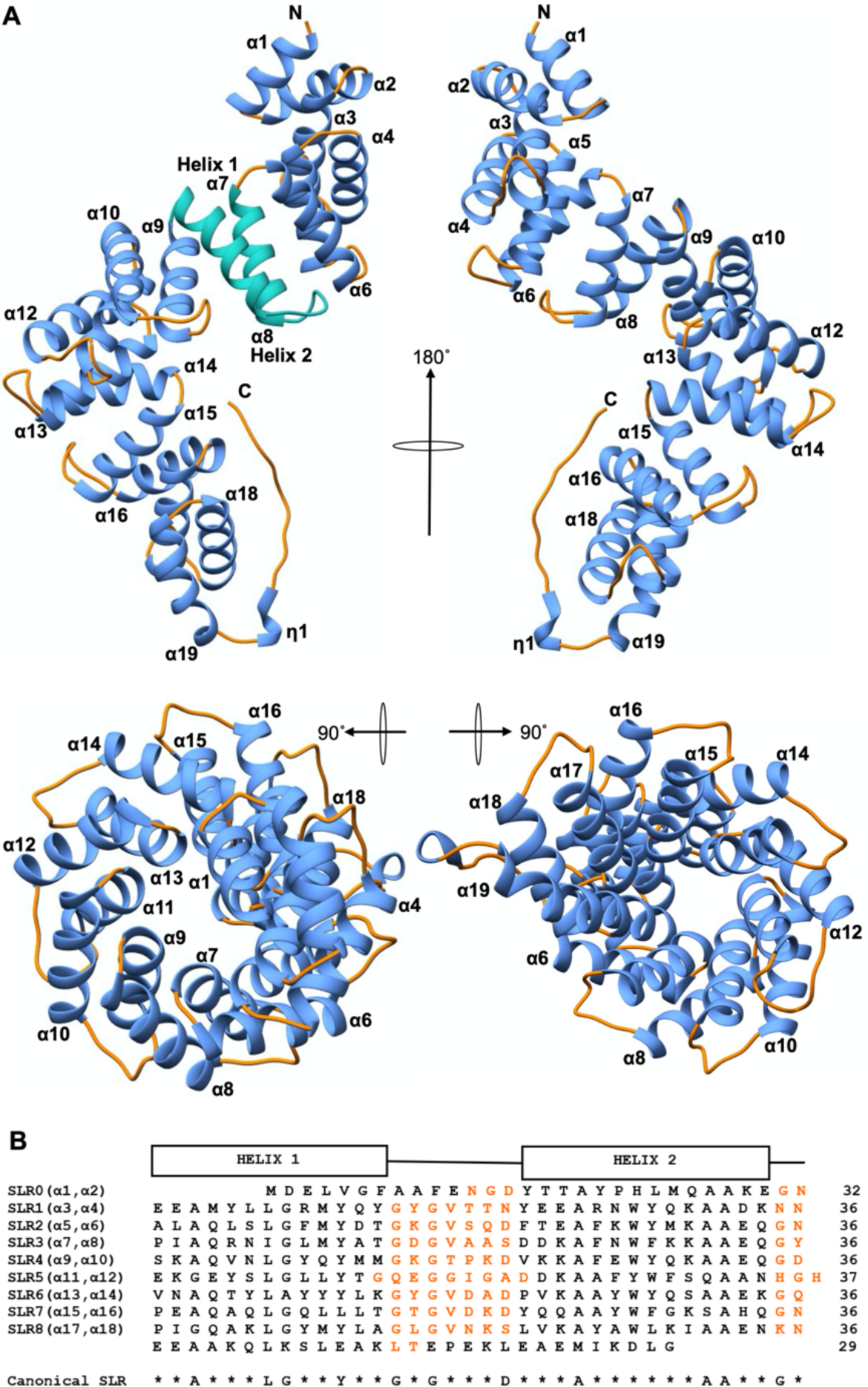
X-ray crystal structure of LceB. A) four views of *Legionella pneumophila* LceB are shown as a ribbon diagram. The N and C terminus are labeled with secondary structure indicated as alpha-helices (α) and 3_10_ helices as (η). Helices are colored in *blue* or *green* and loop regions in *orange*. A single SLR made up of Helix 1 (α7) and Helix 2 (α8) is depicted in *sea green*. B) Alignment of LceB SLR’s. Secondary structure elements can be observed above the SLR sequences (Helix 1 and 2). The canonical SLR is located below the sequences with loop regions colored *orange*.

**Table 1.** X-ray data collection and refinement statistics.

| Data Collection | LceB |
| --- | --- |
| Wavelength (Å) | 1.18082 |
| Space Group | P3<sub>2</sub> |
| Cell dimensions |  |
| a, b, c (Å) | 116.79, 116.79, 88.35 |
| α, β, γ (°) | 90.0, 90.0, 120.0 |
| Number of monomers in asymmetric unit | 3 |
| Resolution (Å) | 48.72 – 2.70 (2.80-2.70) |
| Total reflections | 190820 (23457) |
| Unique reflections | 36973 (3696) |
| CC(1/2) | 0.999 (0.499) |
| R<sub>merge</sub> | 0.021 (1.159) |
| R<sub>pim</sub> | 0.011 (0.565) |
| I/σI | 20.5 (2.69) |
| Completeness (%) | 99.91 (100.0) |
| Multiplicity | 4.7 (5.2) |
| Refinement |  |
| R<sub>work</sub>/R<sub>free</sub> | 0.2169/0.2509 |
| Average B-factors (Å<sup>2</sup>) | 111.35 |
| Protein | 111.51 |
| Ligands | 99.98 |
| Water | 89.89 |
| No. atoms | 8279 |
| Protein | 8207 |
| Ligands | 66 |
| Water | 46 |
| Rms deviations |  |
| Bond lengths (Å) | 0.002 |
| Bond angles (°) | 0.38 |
| Ramachandran plot (%) |  |
| Total favoured | 93.67 |
| Total allowed | 6.14 |
| PDB code | 8SXQ |

As shown in Figure 1, LceB is composed primarily of α-helices forming a right-handed superhelix that is ∼93 Å long ending in an extended C-terminal loop region. There are 19 α-helices which form eight pairs of antiparallel helices with each pair making up a Sel1-like repeat (SLR). A single Sel1-like repeat (SLR3) is highlighted in turquoise for clarity as helix 1 (α7) and helix 2 (α8) in Figure 1A with the sequences of each repeat unit shown in Figure 1B. The concave outer surface of the LceB superhelix is composed of Helix 1 and the convex inner surface is made of Helix 2 of each SLR. Of note, is that the first two α-helices (α1 and α2) at the N-terminus of LceB are not composed of the required number of residues or adhere to the canonical SLR sequence. However, these N-terminal helices conform to the SLR or TPR fold. Additionally, the C-terminal helix (α19) appears to have no mate and we hypothesize that it acts as a capping helix to shelter the concave hydrophobic regions which are found in solenoid proteins like LceB^34^. For example, in TPR proteins, capping helices are common and may help to increase protein stability and solubility^29^. Finally, LceB terminates in a 3_10_ helix that leads into a long-extended tail.

The SLRs in LceB are separated from each other by two residues except for SLR5 and SLR6, where three residues make up the loop region (Fig. 1B). Typically, the first residue of these loops are glycines, and are overall strictly conserved while the second residue is variable. Additionally, all the SLR inter-helix loop regions in LceB are made up of seven residues except for SLR5 which contains nine. It is important to note that two definitions exist for describing SLR proteins^20, 46^.. The definition used in this paper is that from the Smart database^20^, where the length of the loop regions between helices within one SLR are long (4-12 residues) and the connecting regions between SLRs are short (3 residues)^34^. This was chosen as LceB conforms to the consensus sequence of SLRs described by the Smart database and not the other SLR description. Interestingly, the alternative definition is used in the structural characterization of LpnE in *L. pneumophila*^47^_._

### Molecular surface properties of LceB

To gain insight into a biochemical function for LceB, we first analyzed the structure for surface residue conservation for a potential active site or binding-partner surface. As shown in Figure 2A, the majority of the LceB surface shows no residue conservation. Notably, the areas of most variability are at the N and C-termini. This includes α1 and α2 that conform to the SLR fold but do not contain the SLR sequence, and the capping helix α19 with the extended C-terminal tail. Additionally, no areas of residue conservation or predicted active site motifs^48^ are readily apparent on the concave surface of LceB. However, there are repeating patches of conservation along the entire length of the convex surface and defined stripes (Fig. 2A). This contrasts with TPR proteins where the concave surface is commonly responsible for binding interactions^29^.

**Figure 2.**
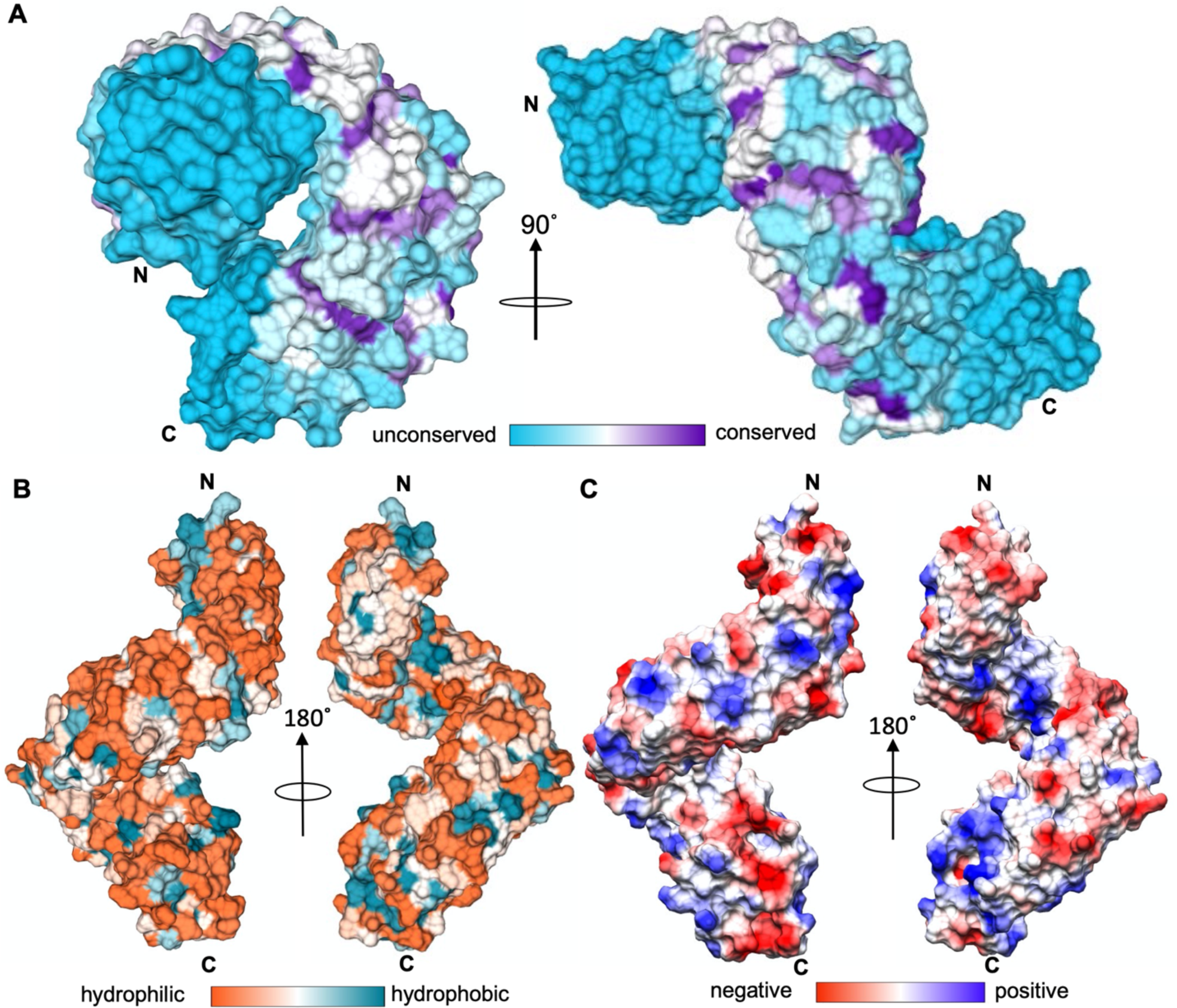
Molecular surface properties of LceB. A) Residue surface conservation of LceB generated by Consurf (https://consurf.tau.ac.il/). Unconserved surfaces are colored in *turquoise* and conserved surfaces in *deep purple*. Areas of *white* or light shades indicate partial conservation or homologous residue regions. B) Hydrophobic surface representation of LceB. A gradient of hydrophilic surfaces (*orange*) to hydrophobic patches (*blue-green*) is drawn. C) LceB colored by Coulombic surface or electrostatic potential ranging from negative surface (*red*) to a positive surface (*blue*) as calculated by ChimeraX. In each panel the N and C termini of LceB are indicated.

We next plotted both the hydrophobic and electrostatic properties of the LceB surface to observe if any of these properties align to the repeating patches of conservation in LceB (Fig. 2B and Fig. 2C). As shown, the surface of LceB is very hydrophilic with few exposed non-polar residues and no obvious hydrophobic patches (Fig. 2B). The N-terminal surface of LceB shows a slight overall negative charge while the C-terminal convex region appears to have a positively charged patch (Fig. 2C). As both the N and C-terminal regions show low residue conservation, it is unlikely that these electrostatics correlate with a conserved SLR-domain function. Overall, neither hydrophobic regions or electrostatic regions aligned to areas of residue conservation in LceB (Fig. 2). Regardless, as TPR family proteins are variable in sequence due to their unique functions^49, 50^ these observed molecular features could be important for biological partner interactions specific to LceB.

### LceB contains motifs of known function

LceB exhibits a repeated motif (KAAEQG) and is always within the second helix of an SLR. The KAAEQG motif can be observed in SLR2 (α5-6), SLR3 (α7-8), and SLR4 (α9-10) and represents a partially conserved surface (pink) on LceB (Fig. 1B and Fig. 3A). Importantly, the motif itself is highly conserved especially at the second alanine and terminal glycine residue. Substitution of other positions are always with a homologous residue. SLR1 (α3-4), SLR5 (α11-12), and SLR6 (α13-14) contain variations to the KAAEQG motif and bound both sides of the conserved KAAEQG surface (brown, Fig. 3A). Namely, SLR1 (α3-4) has the terminal glycine replaced with lysine and both SLR5 (α11-12), and SLR6 (α13-14) have the initial lysine substituted to serine (Fig. 3B). A search for LceB structural homologs using the Dali server^51^ showed that the KAAEQG sequence and its variations are found in all the listed LceB structural relatives except for HcpC (Table 2). In many SLR proteins, and observed in LceB, the conserved alanine and glycine residues allow tight packing of the repeats^41^. Additionally, this motif pattern is considered to be important to maintain the angular geometry between repeats, and thus the overall structure of the SLR fold^41^. However, as the KAAEQG and variant motifs also create a partially conserved surface found on the edge of the convex surface of LceB, this may indicate a potential binding site for a biological interaction partner.

**Figure 3.**
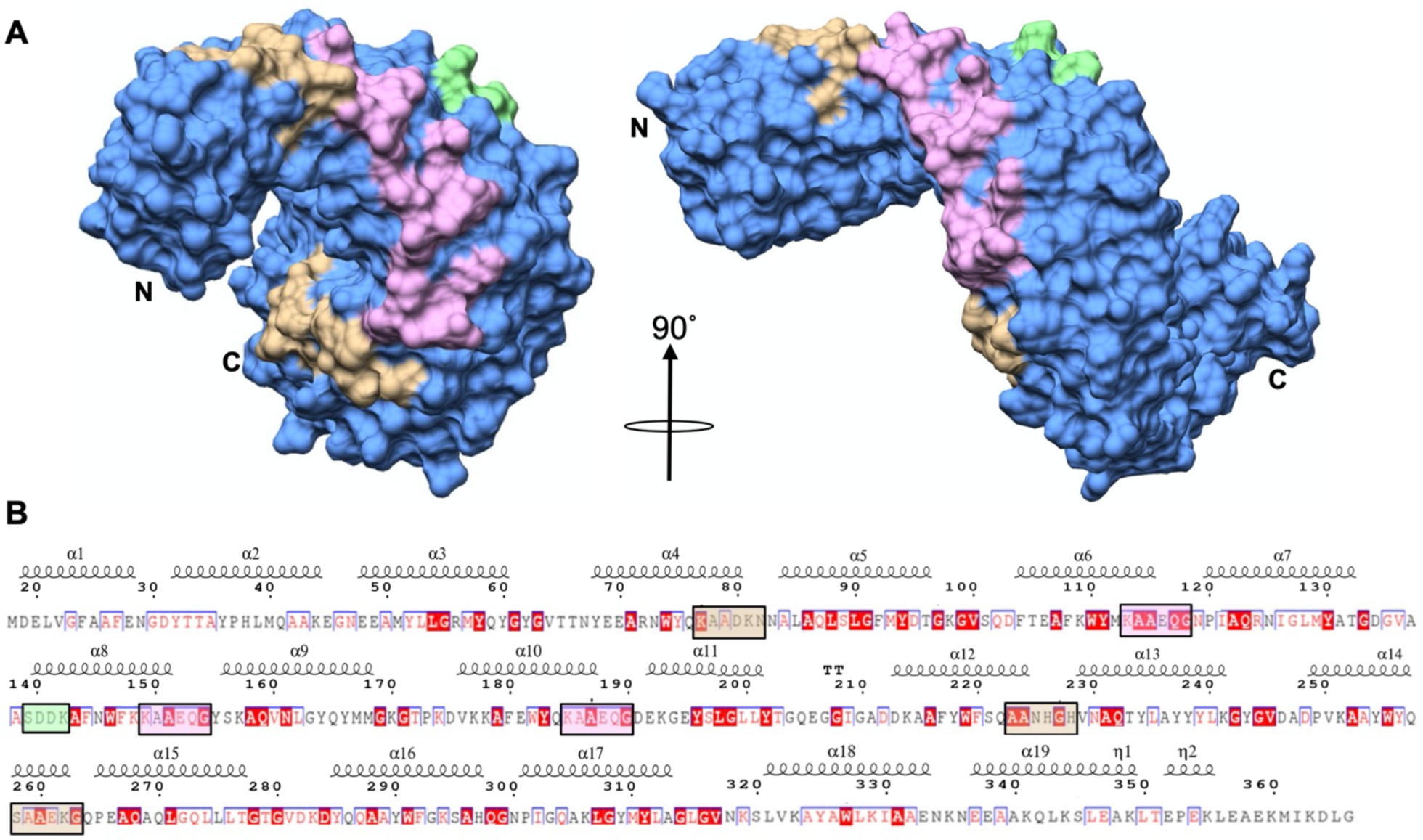
Conserved surface residue motifs of LceB. A) Molecular surface representation of LceB showing predicted functional regions of LceB. The repeated KAAEQG sequence motif is highlighted in *pink* and is bounded by two closely related motifs in *brown*. A catalytic serine motif (SXXK) observed in known SLR β-lactamases is colored green. B) LceB amino acid sequence plotted with corresponding secondary structure and residue conservation using Espript (https://espript.ibcp.fr/). Residues highlighted in *red* indicate strict conservation, homologous residues are colored *red*, while unconserved residues are colored *black*.

**Table 2.**
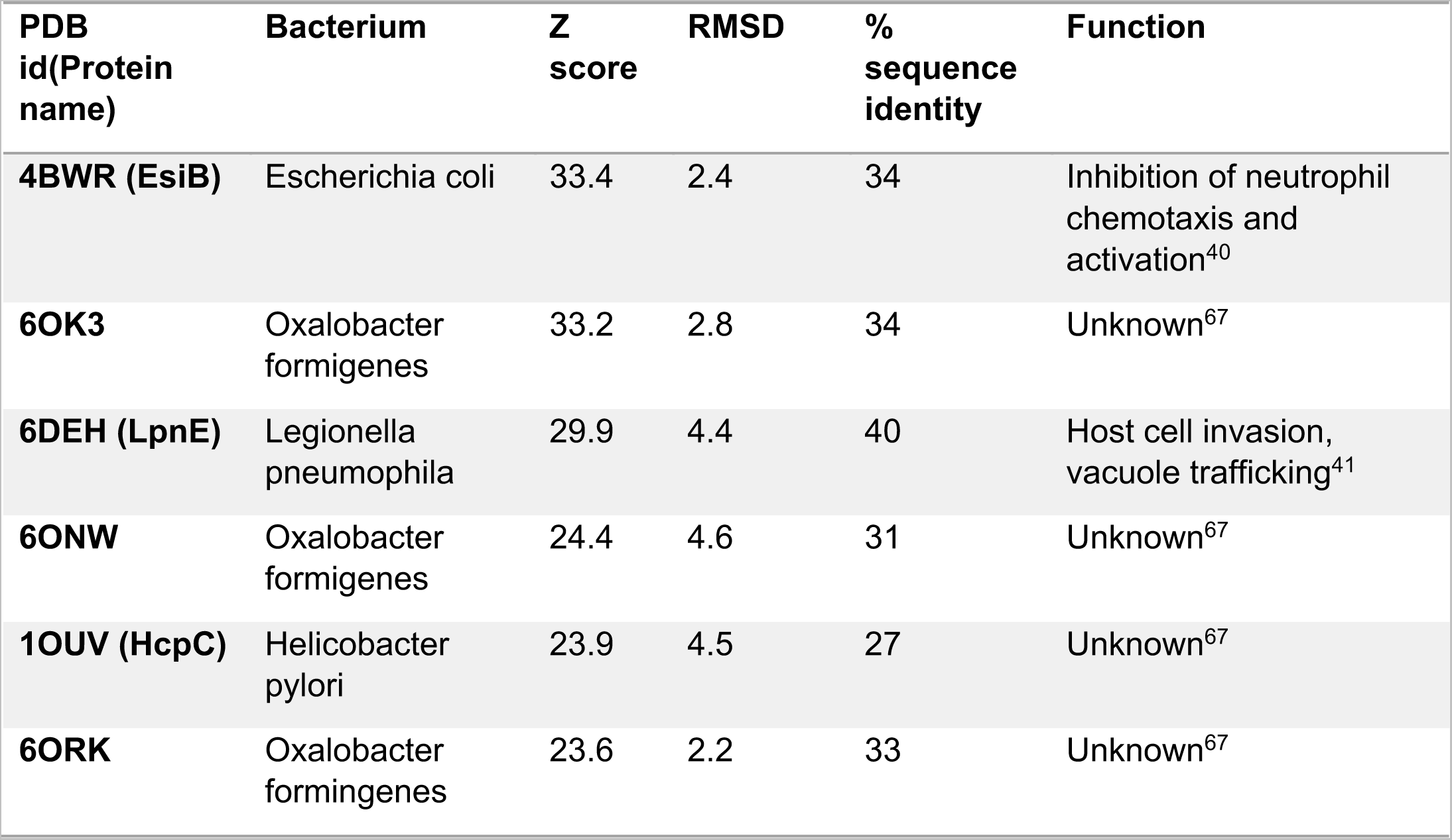
LceB structural homologs.

| **PDB id(Protein name)** | **Bacterium** | **Z score** | **RMSD** | **% sequence identity** | **Function** |
| --- | --- | --- | --- | --- | --- |
| **4BWR (EsiB)** | Escherichia coli | 33.4 | 2.4 | 34 | Inhibition of neutrophil chemotaxis and activation<sup>40</sup> |
| **6OK3** | Oxalobacter formigenes | 33.2 | 2.8 | 34 | Unknown<sup>67</sup> |
| **6DEH (LpnE)** | Legionella pneumophila | 29.9 | 4.4 | 40 | Host cell invasion, vacuole trafficking<sup>41</sup> |
| **6ONW** | Oxalobacter formigenes | 24.4 | 4.6 | 31 | Unknown<sup>67</sup> |
| **1OUV (HcpC)** | Helicobacter pylori | 23.9 | 4.5 | 27 | Unknown<sup>67</sup> |
| **6ORK** | Oxalobacter formigenes | 23.6 | 2.2 | 33 | Unknown<sup>67</sup> |

LceB also contains an SXXK motif on its surface depicted in green (Fig. 3A). This motif is found in the structurally homologous protein HcpC from *H. pylori* (Table 2). HcpC is from a family of *Helicobacter* cysteine-rich proteins (Hcps) found in epsilon proteobacteria^46^ which includes HcpA, B, D and E also found in *H. pylori*^34, 46^. Members of this family of proteins have been shown to bind and weakly hydrolyze derivatives of penicillin^33, 52^. The active site in β-lactamases contain a catalytic serine within a SXXK motif where the lysine is invariable and two other motifs, ((S/Y)X(N/S/D/C)) and (K/L)(T/S)G) separated by approximately 70 residues^59, 60^. Interestingly, HcpA contains only the SXXK and ((S/Y)X(N/S/D/C)) motifs yet retains hydrolysis activity^52^. Additionally, HcpB was shown to co-purify with N-acetylmuramic acid and has all three motifs^33^. Overall, these results suggested that Hcps may be involved in cell-wall biosynthesis^33, 46^. Given that LceB has a sec-signal and contains a SXXK motif, it is tempting to speculate that it may have a dual function in the periplasm or is also part of the T4SS apparatus.

The closest known structural homolog to LceB is the protein EsiB (Table 2 and Fig. 4A). As shown in Figure 4A, LceB aligns extremely well to the core of EsiB but with variations at each terminus. Namely, EsiB has 3 additional Sel1-like repeats at the N-terminus and the packing of the C-terminal domain varies. Similar to LceB, EsiB also contains a capping helix at the C-terminus and follows the same Sel1-like repeat definition as LceB. Again, this definition is that each SLR consists of two α-helices linked by 7 amino acids with three residues connecting adjacent SLRs^41^. EsiB is a virulence factor found in uropathogenic *Escherichia coli* (UPEC), a subtype of extraintestinal pathogenic *E.coli* (ExPEC), CFT073 *Escherichia coli*^41^. ExPEC is pathogenic to both humans and animals causing neonatal meningitis (NMEC), septicemia as well as urinary tract infections (UTIs)^54^. Mucosal surfaces in humans are protected considerably by secretory immunoglobulin A (SIgA) which is hydrophilic and negatively charged^55^. SIgA also binds to the FcαRI receptor on neutrophils leading to immune cell activation and thus destruction of the pathogen^55^. EsiB functions by binding to SIgA without blocking its interaction with the FcαRI receptor. Overall, this results in the failure of the neutrophil to be activated, aiding in the survival of the pathogen^55^. Moreover, the biochemical activity of EsiB hinders chemotaxis of neutrophils^55^. The specific amino acid sequence in EsiB that interacts with SlgA is 244-VLFS<u>QSAEQG</u>NSIAQFR-260^41^. Variations of this sequence can be observed throughout both EsiB and LceB (Fig. 3B, Fig. 4B). Specifically, the SlgA binding motif includes the KAAEQG repeat motif of LceB (underlined in the SlgA binding sequence). This close structural and sequence motif identity of LceB to EsiB could indicate that the role of LceB in the LCV is to inhibit an immune response.

**Figure 4.**
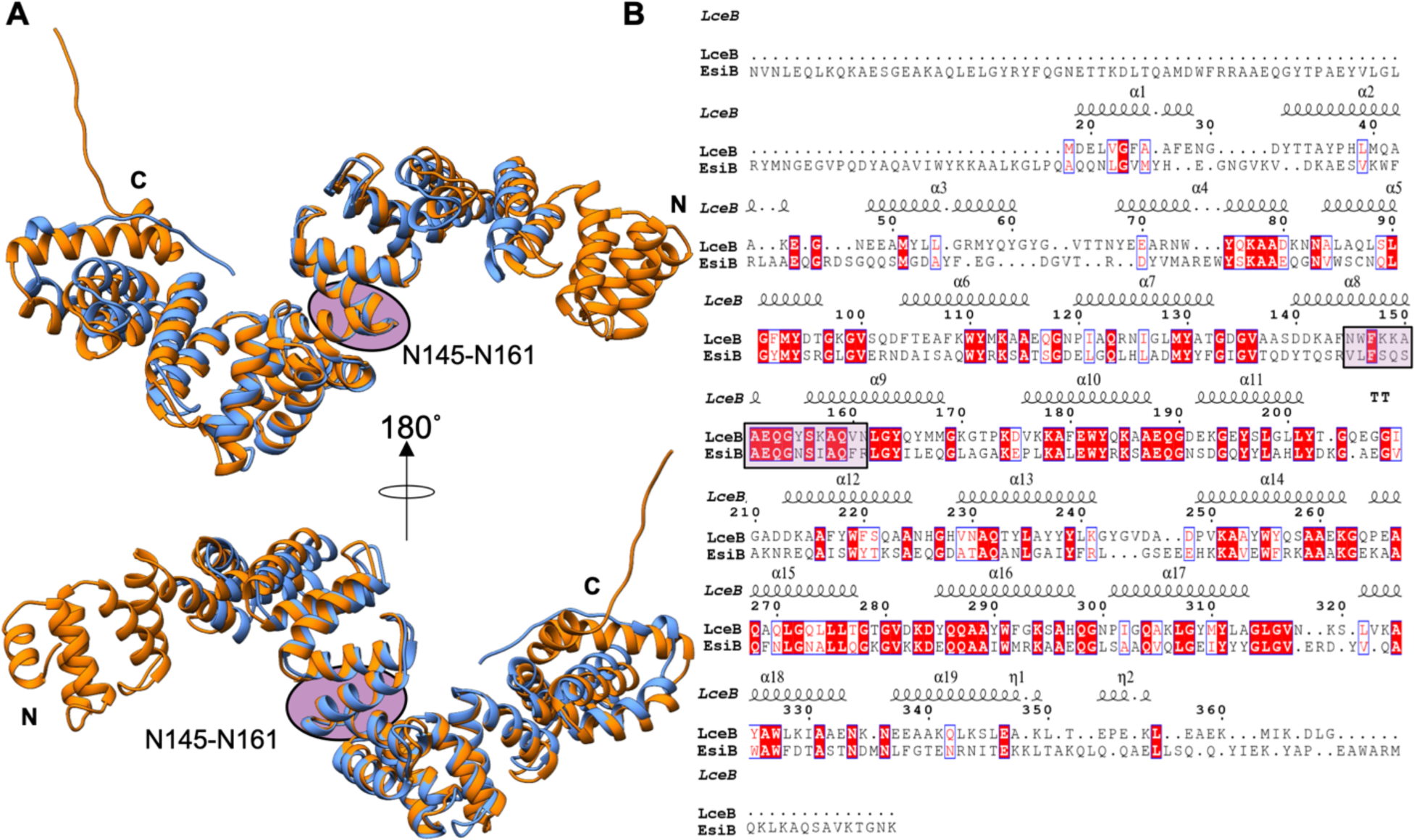
Structural alignment of LceB with EsiB from *E. coli*. A) LceB (*cornflower blue*) aligned with EsiB (PDB 4BWR) (*orange*) is shown in two views with a 180° rotation. B) The sequence alignment based on the structural alignment of LceB and EsiB from the Daliserver is shown. The secondary structural elements and residue conservation is plotted by Espript (https://espript.ibcp.fr/). Residues highlighted in *red* indicate strict conservation, homologous residues are colored *red*, while unconserved residues are not bolded and colored *black*. The sequence that EsiB uses to bind SlgA is highlighted with a *purple* oval on the structures in panel A and by a box on the sequence in panel B. This corresponds to a KAAEQG sequence motif.

### Structural Comparison of LceB to known SLR proteins

In addition to HcpC and EsiB, LceB has several additional structural homologs as determined by the DALI server^51^. The top six non-redundant structural hits had Z-scores ranging from 33.4 to 23.6 with the remaining hits all scoring below a Z-score of 20 (Table 2). The RMSD and sequence identity for the top six hits ranged from 2.2 Å^2^ to 4.6 Å^2^ and 27% to 40%, respectively. Intriguingly, three of the six hits (PDBid: 6OK3, 6ONW, 6ORK) are from *Oxalobacter formigenes*, a major degrader of oxalate in the human gut^56^, but the function of each of these proteins has not yet been determined. However, the predicted annotation for each of these gene products is beta-lactamase and carbon-nitrogen bond hydrolase activity.

LceB is also structurally related to LpnE (PDB 6DEH) from *L. pneumophila*^51^. LpnE is a well characterized SLR family protein that has both known protein and lipid binding partners. LpnE binds Obscurin-like protein 1 (OBSL1) directly at the OBSL1 eukaryotic immunoglobulin-type folds *in vitro*, phosphatidylinositol-3-phosphate (PI3P) and Oculocerebrorenal syndrome of Lowe protein (OCRL)^42, 57^. These binding activities occur within the LCV during host invasion^42, 57^. OCRL dephosphorylates phosphatidylinositol-4,5-bisphosphate (PtdIns(4,5)P_2_) and has been shown to impede *Legionella* infection^57–59^. LpnE has previously been shown to be important in trafficking of vacuoles and host cell invasion, an activity that appears to require all eight of its SLR motifs^42^. It is important to note that two of the other 5 known *Legionella* SLR proteins LidL and EnhC are also involved in vacuolar trafficking^39, 40, 42, 43^. For example, EnhC is able to complement an LpnE mutant strain, and LpnE is able to complement an EnhC deficient strain^60^. This common phenotype, coupled with the fact that Dot/Icm effectors are redundant^61^ further suggests that LceB may aid in maintaining the LCV within human macrophages.

Several general structural similarities also exist when LceB is compared to other SLR proteins. Like LceB, many SLR proteins contain secretion and signal sequences (EsiB, EnhC, HcpC, LpnE). For example, the N-terminal signal sequence (1-21) of LpnE is responsible for localization to the *cis*-Golgi in HEK293 cells and without the sequence LpnE is found to be retained within the host cytosol^47^. Given this, it is likely that LceB requires a specific localization to exert its biological function. The LceB structural homologs span a repeat range of 7 to 21 SLRs and homologues such as EsiB, HcpC, and LpnE also exhibit a C-terminal capping helix that serves to stabilize the overall curved SLR structure^41, 42, 46^. Furthermore, many residues are conserved for structural purposes in the SLR proteins. Alanines in positions 3 and 32 (LceB numbering) are highly conserved within each SLR which allows tight packing of the repeats^41^ (Fig. 1B and Fig. 3B). Specifically, the conserved alanine at the C-terminus of an SLR associates with another conserved alanine at the N-terminal of a subsequent repeat to properly pack the repeats together. Additionally, the tryptophan at position 27 (LceB numbering) although not part of the canonical SLR sequence is conserved in nearly all the Sel1-like repeat homologs listed in Table 2 and makes stabilizing contacts with the previous SLR in the sequence. The pattern for the residue associations are also seen in EsiB where the authors hypothesized that this interaction was important for maintaining the angular geometry between repeats^41^.

### LceB shows structural flexibility

The three chains of LceB in the asymmetric unit (Fig. S2) were captured in different conformations within the crystal (Fig. 5A). Chain A superimposes on Chain B with an RMSD of 0.8 Å^2^ across 198 equivalent C-alpha atoms (2.6 Å^2^ for all 348 residues) and Chain C of 0.9 Å^2^ for 325 Cα atoms (1.2Å^2^ for all). Chain B and C superimpose with an RMSD of 0.9 Å^2^ (168 Cα atoms) and 3.4Å^2^ for all atoms. Between the three chains, the greatest variation was seen in both the N-terminal (SLRs 0 to 2 or helices α1-α6) and C-terminal regions (SLR 8 or helices α17-α18). This includes the C-terminal residues that appear to adopt a poorly resolved helix that ends in an extended conformation (Fig. S3). Overall, it appears that LceB can ‘flex’ to further open the inner concave surface as the C-terminus of Chain B is pushed ∼ 6Å out relative to Chain A (Fig. 5A).

**Figure 5.**
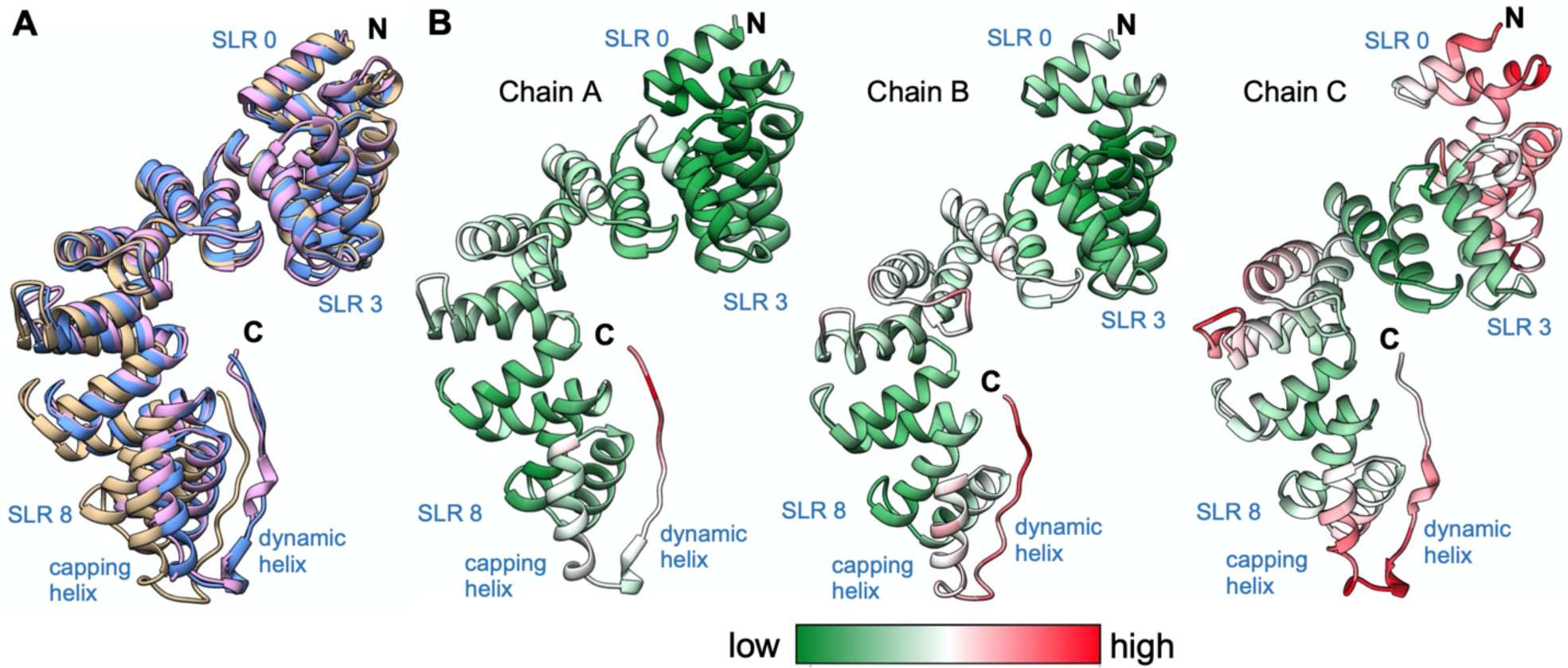
Comparison of the three LceB copies in the asymmetric unit. A) Superimposition of LceB chains. Chain A is in *cornflower blue*, Chain B is in *tan*, and Chain C is in *plum*. All panels have the N and C termini labeled. B) For each chain the observed B-factors are colored by gradient from low (*green*) to red (*high*). Each chain has the N and C terminus indicated with relevant structural features highlighted.

Plotting the B-factors for each chain in the LceB crystal structure reveals varying levels of motion across the protein. Chain A showed the least motion followed by chain B with chain C having the highest overall thermal parameters (Fig. 5B). In chain C, both the N and C-terminal regions had the highest observed B-factors suggesting these regions of LceB may be flexible relative to the rest of the protein. Although the thermal parameters could relate to crystal packing artifacts, it must be noted that in all three chains the C-terminal region was exposed to solvent and had poor electron density (Fig. S3). The density appears to suggest an α-helix, and in fact, the AlphaFold model used for molecular replacement predicts the C-terminus as a long α-helix (Fig. 6A). Taken together, the experimental data and the predictive model indicate that the C-terminal region of LceB is likely a conformationally dynamic helix. Given that this last helix is beyond the capping helix (α19), sequence variable, and not a canonical SLR mate, the structural and biochemical role of this region is unclear.

**Figure 6.**
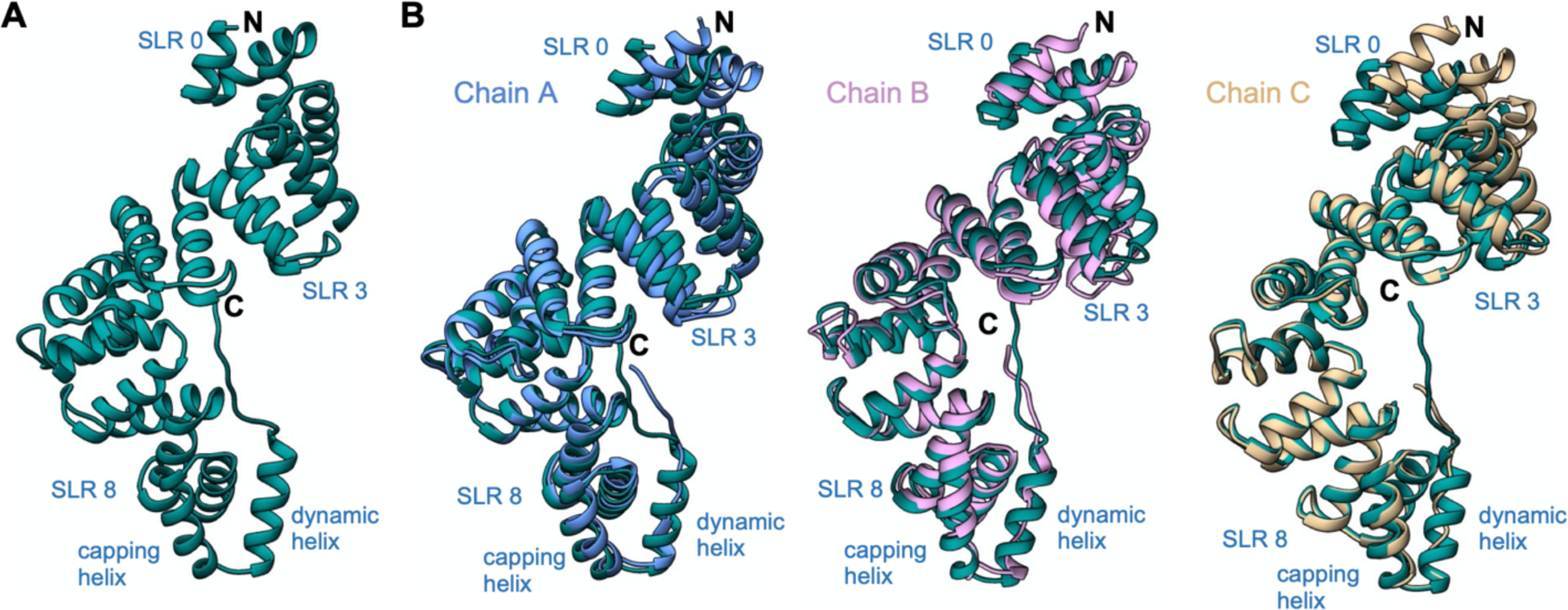
The crystal structure of LceB varies from the AlphaFold model. A) LceB structure predicted by AlphaFold and used as a molecular replacement model. B) The Alphafold model (*steel blue*) aligned with each chain in the LceB X-ray crystal structure. chain A (*cornflower blue*), chain B (*plum*), and chain C (*tan*). Each chain has the N and C terminus indicated with relevant structural features highlighted.

Further supporting the experimental data that LceB is conformationally dynamic, the AlphaFold model predicts a different overall conformation for LceB as compared to the X-ray crystal structure. In addition to showing a folded C-terminal helix, the packing arrangement of SLRs 0 to 3 in the AlphaFold model is bent so as to close the concave surface (Fig 5B). For both chain A and chain C, SLR 0 (α1 and α2) is moved about 4.5Å outwards relative to the predicted structure with differences in the position of helices averaging ∼4Å in the entire N-terminal half of LceB. The conformational change is even more pronounced when compared to chain B. Helices α1-α5, or SLRs 0-1 and half of SLR 2, in the experimental structure are more than 7Å and up to just over ∼8Å shifted outward relative to the AlphaFold model. When comparing the predicted model and the three LceB chains of the experimental structure, we observe a closed conformation in the AlphaFold model and an open conformation in chain B, with both chain A and C roughly at the midpoint between the extremes.

### LceB oligomerizes in solution

LceB was readily purified by metal chelating affinity chromatography followed by size exclusion chromatography (SEC) where it was observed to elute as two distinct peaks (Fig. 7A). Although no obvious dimer or larger oligomer was apparent from the asymmetric unit in the crystal structure (Fig. S2), this suggests that LceB may be in equilibrium between a monomer and higher order oligomer. Alternatively, the two peaks could represent a compact and elongated conformation of LceB as suggested by the observed flexibility and dynamic C-terminus (Fig. 5 and Fig. 6).

**Figure 7.**
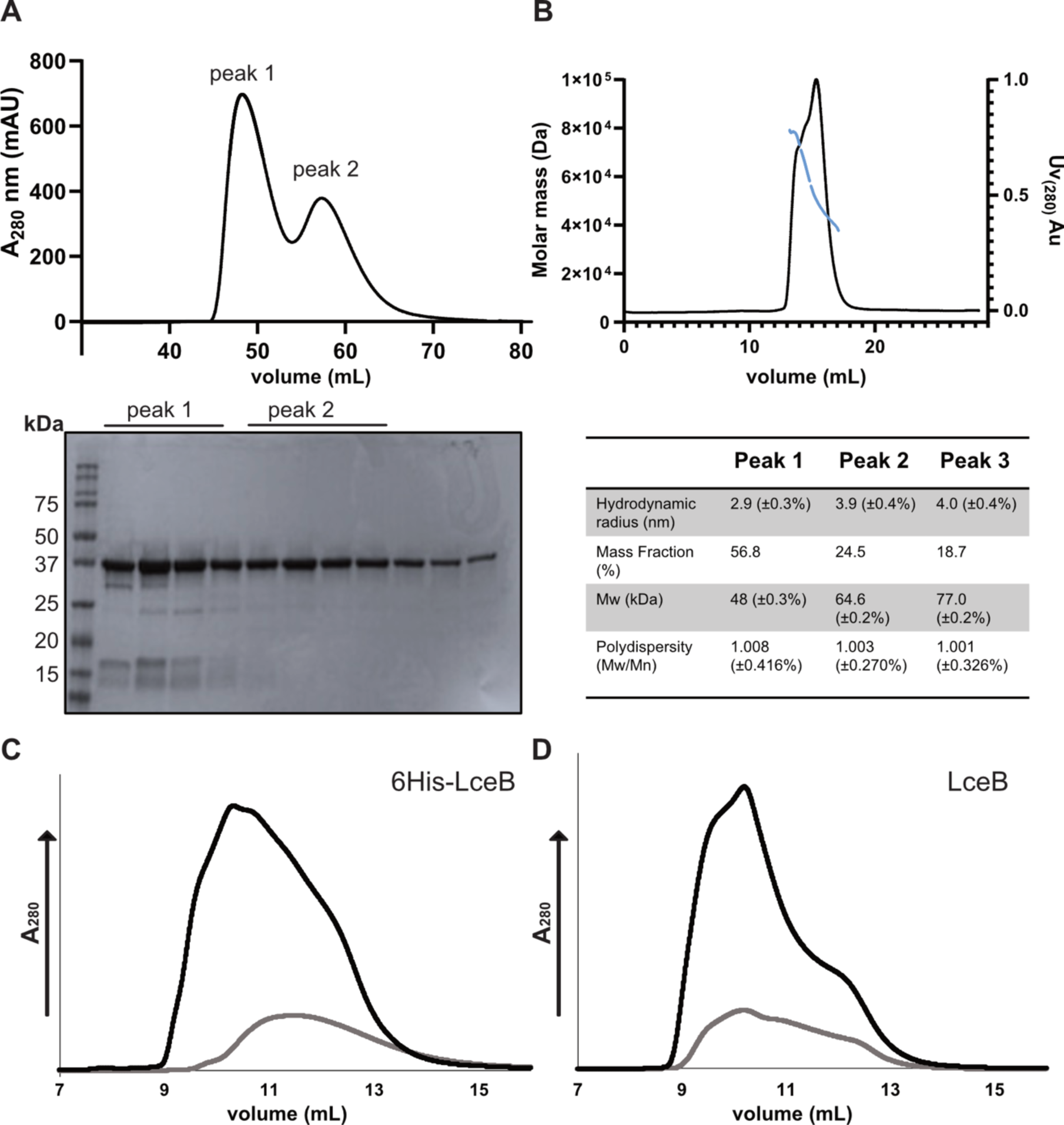
Solution state analysis of LceB. A) SEC trace of 6His-LceB after nickel affinity chromatography (top) and Coomassie stained SDS-PAGE gel of purified 6His-LceB SEC fractions (bottom). B) SEC-MALS trace of peak 2 from panel A showing calculated molecular mass fit of the peak(s) in *blue* (top) and mass-analysis from ASTRA analysis software (bottom). C) SEC trace of 6His-LceB at 5 mg/mL (*black*) and 1 mg/mL (*grey*). D) SEC trace of LceB after 6His-tag removal at 5 mg/mL (*black*) and 1 mg/mL (*grey*). Graphs were plotted using GraphPad Prism version 9.4.1.

To better determine the oligomeric state of LceB, the fractions from peak 2 (Fig. 7A) were concentrated and analyzed by size exclusion chromatography coupled with multi-angle light scattering (SEC-MALS). In this experiment, LceB eluted as three peaks. Peak one yielded a measured mass of 48 kDa and made up more than half of the mass fraction (56.8%) (Fig. 7B). This peak likely corresponds to a monomeric species as the predicted molecular mass of an LceB monomer is 42 kDa. Peaks two and three yielded molecular masses and mass fractions of 64.6 kDa (24.5%) and 77 kDa (18.7%) respectively. Neither of these two measured molecular masses are enough to match that of a dimer (96 kDa) in this experiment, and together compose less than half of the protein sample. By looking at the hydrodynamic radius value of the monomer (2.9 nm) and comparing it to peak 2 and 3, (3.9 nm and 4.0 nm respectively), it is possible that these peaks show LceB monomers associated with two different confirmations or are elongated forms of the protein.

In agreement with our observations, a study by Voth and colleagues in 2019 investigating the LceB homolog LpnE found similar behavior in solution^47^. When SEC was performed with His-tagged LpnE, the protein also eluted in two peaks. However, when SEC-MALS was used to study an untagged version of LpnE, the protein eluted as a monomer. This suggested that the His-tag was responsible for the self-association or dimerization of LpnE^47^. Given this similarity, we expect the larger particle size peaks observed for LceB to also be due to cloning artifacts. To address this possibility, we removed the N-terminal 6His-tag by digestion with thrombin (Fig. S4) and performed SEC analysis of 6His-LceB and digested LceB at different concentrations (Fig. 7C and Fig. 7D). As shown in the chromatograms neither concentration nor the removal of the affinity tag resulted in uniform peak, or single species in solution. Either with or without the 6His-tag, LceB appears to aggregate in solution. However, without the affinity tag LceB seems to favor either an extended conformation or larger order oligomer in solution. This can be observed as 6His-LceB shows more material at an elution volume of 11 mL or higher, whereas digested LceB has a significant peak at ∼9.8 mL regardless of concentration (Fig. 7C and Fig. 7D). This data indicates that LceB does self-interact independent of cloning artifacts, however the exact nature of the LceB oligomer is unknown.

To determine the exact nature of the LceB oligomers, we measured the oligomeric state of Lpg1356 in solution by mass photometry. This analysis revealed three major peaks corresponding to the estimated MW of a monomer (peak 1), trimer (peak 2), and pentamer (peak 3) of LceB (Figure 8). Oligomer sizes were estimated from the monomer appearing to be 51 kDa in solution. These states accounted for over 80% of the analyzed sample. Additionally, minor species corresponding to a decamer (peak 4) and possible 12 to 14-mer (peak 5) of LceB were detected. Given this behavior, it is tempting to speculate that the 3 copies of LceB in the asymmetric unit may be reflective of the solution-state trimer, or at least provides insight into how LceB may self-associate (Fig. S2). However, this seems unlikely as none of the crystal-contacts contained conserved surface residues.

**Figure 8.**
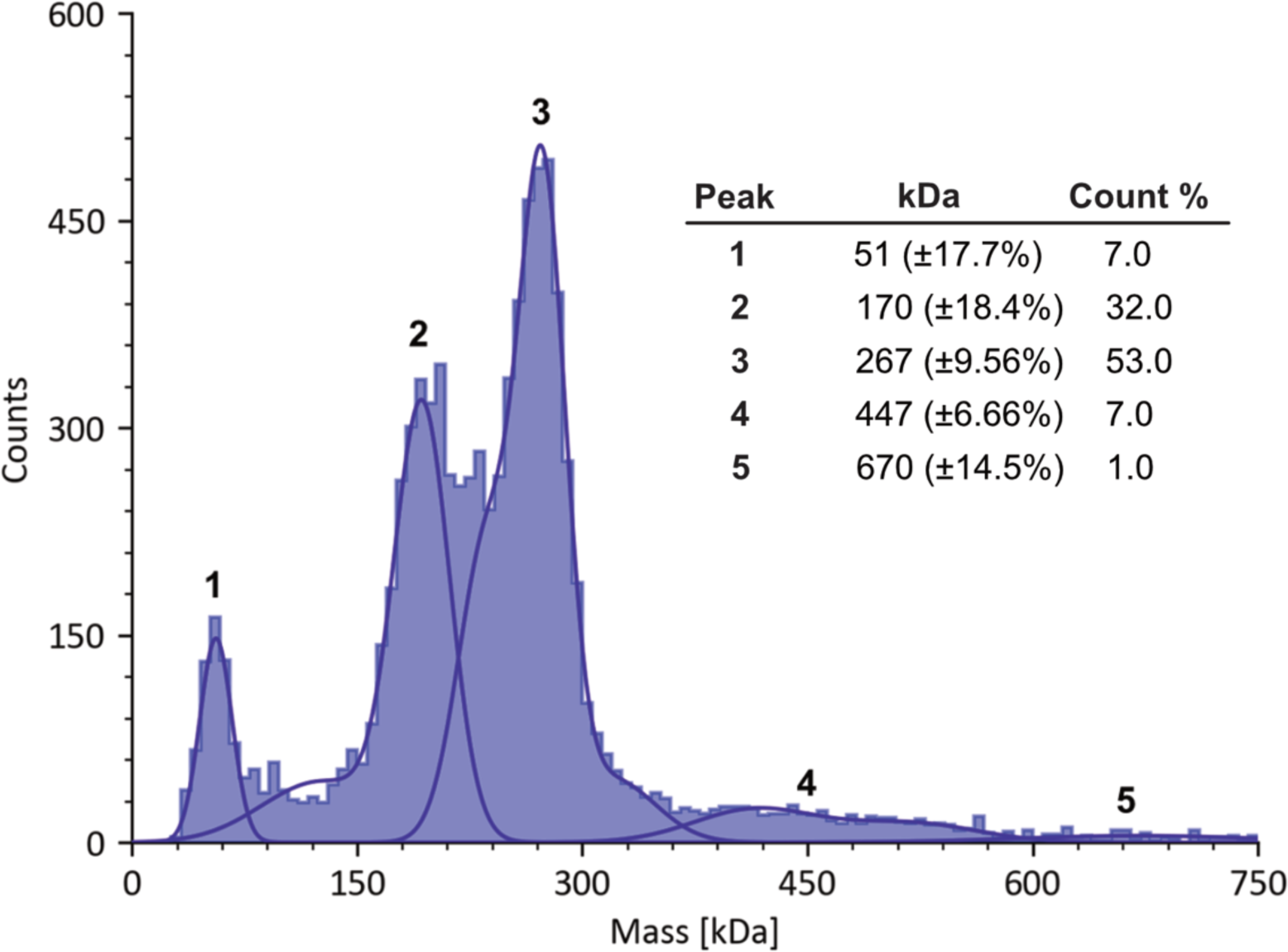
LceB forms several higher-order oligomers in solution. Molecular weight histogram obtained from Mass photometry measurements of purified LceB. Approximate oligomeric states are Peak 1 = monomer, Peak 2 = trimer, Peak 3 = pentamer or hexamer, Peak 4 = decamer, and Peak 5 = unknown multimer. Measurements were repeated three times and produced similar results.

## CONCLUSION

An X-ray crystal structure of LceB from the respiratory pathogen *Legionella pneumophila* has been solved to a resolution of 2.7Å in space group P3_2._ LceB crystallized as 3 super-helical chains in the asymmetric unit, with each chain composed of 8 Sel1-like repeats (SLRs) (Fig. 1). Upon structural analysis we observed that LceB contains a repeated motif, KAAEQG that is also found in its closest structural homologs (Fig. 3). Although this motif has a structural role^41^, it may also be functionally important. Namely, the closest structural homolog to LceB, the protein EsiB in uropathogenic *E. coli* requires this motif to bind the host protein SIgA (Fig. 4 and Table 2). The binding of EsiB to SlgA interferes with the host response inhibiting neutrophil activation, leading to pathogen survival^55^. It is therefore likely that this repeated conserved motif sequence in LceB is also the site of protein binding, especially as it forms a conserved surface on LceB (Fig. 2A and Fig. 3A). However, a protein binding partner for LceB has yet to be determined. Additionally, LceB is structurally similar to another *Legionella* SLR protein, LpnE, which functions in vacuolar trafficking during host cell invasion. Taken together, this may be suggestive that LceB has multiple functions. Interestingly, both these proteins have a general secretory signal which at least for LpnE, is thought to be necessary for its localization within a host. This raises the question of why LceB contains a sec-signal, especially as data exists that it is secreted by the sec-independent T4SS^17^.

Our structural data also indicates that LceB is conformationally dynamic, or at the very least flexible in solution. The chains in the asymmetric unit showed different conformations, with chain B showing the most variation from chain A and chain C (Fig. 5A). Furthermore, the AlphaFold model of LceB also predicted a significantly different conformation from those observed in the crystal structure, especially in the N-terminus the protein (Fig. 6). This, combined with the high B-factors and poor density for the putative C-terminal helix, strongly indicates that LceB is conformationally dynamic in solution. Moreover, LceB readily forms trimers and higher-order oligomers in solution (Fig. 7 and Fig. 8) which could indicate it has adhesin-like biological properties to form large complexes with its eukaryotic host target. Overall, our determination of an experimental LceB structure has allowed the exploration of its molecular surface properties and demonstrated a level of conformational flexibility that may shed light not only into its biological function, but also into the structure and biochemistry of the SLR family of proteins. The molecular data of LceB presented here will lead to a better understanding of *Legionella* effector activities and host pathogen interactions.

## MATERIALS AND METHODS

### Protein expression and purification

Codon optimized LceB (*Legionella pneumophila* Philadelphia-1) for *E. coli* was obtained in pET22b vector from Genscript. LceB without the secretion signal was PCR amplified off the plasmid to include restriction sites NdeI and BamHI, and subcloned into pET15b-Amp to generate LceB with an N-terminal 6His-tag in *E. coli* DH5α. Once the expression construct was generated, the vector was transformed into *E. coli* BL21 DE3 Gold cells. The cells were then grown in Lysogeny Broth (LB) at 37 °C until an Optical Density (OD) of 0.6 at 600 nm absorbance was reached. IPTG was added to the culture at a final concentration of 1 mM to induce protein expression for ∼20 h at 20 °C. After incubation, the culture was spun down at 4200*g* for 30 min and resuspended in wash buffer (50 mM Tris pH 7.5, 500 mM or 750 mM NaCl, 25 mM imidazole). PMSF and MgCl_2_ were added at final concentrations of 1 mM and 10 mM respectively, along with a small amount of DNase. An Emulsiflex-C3 High Pressure Homogenizer (Avestin) was employed for cell lysis with the lysate subsequently subjected to centrifugation at 17,000*g* for 30 min at 4 °C. The supernatant was added to a wash buffer-equilibrated nickel-NTA affinity gravity flow column (GoldBio) and elution of the protein was achieved using 1 column volume of elution buffer (50 mM Tris pH 7.5, 500 mM or 750 mM NaCl, 500 mM imidazole). The eluate was concentrated using a 10 kDa concentrator (Sigma) and further purified by size exclusion chromatography (SEC) using a Superdex 75 (16/600) HiLoad column (Cytiva) equilibrated in gel filtration buffer (50 mM Tris pH 7.5, 250 mM or 750 mM NaCl). Protein was further purified by ion exchange chromatography (IEX) after nickel-NTA and SEC. LceB in gel filtration buffer (750 mM NaCl) was diluted to 25 mM NaCl and then eluted by increasing NaCl concentration (0 mM to 300 mM) at pH 7.5 50 mM Tris on an anion exchange (Q-sepharose) column (Cytiva). Fractions from SEC and IEX were run on an 8-12% gradient SDS-PAGE and dyed with Coomassie Brilliant Blue to determine fractions which contained pure protein. Selected fractions were concentrated as previously for crystallization experiments.

### Nano-Differential Scanning Fluorimetry (nanoDSF)

Thermal denaturation of LceB was completed using a Prometheus NT.48 (NanoTemper). Purified protein in 50 mM Tris pH 7.5 250 mM NaCl was diluted 7-fold to a final concentration of 50 µM into Tris and HEPES buffer systems and 25-fold to a final concentration of 10 µM in Bis-Tris buffer systems. Both the pH and salt concentrations were varied. Buffer systems used where: HEPES pH 7.5 and 8, Tris pH 7.5 and 8, Bis-Tris pH 6.5 and 7, sodium acetate pH 3.8 and 4.5. Salt concentrations ranged from 0, 50, 100, 200, 400 and 750 mM NaCl. Samples were then subjected to thermal denaturation by heating from 20 °C to 95 °C at a rate of 1.0 °C/min. Fluorescence intensity was monitored at 330 nm and 350 nm after excitation of tryptophan at 280 nm. Melting temperatures (T_m_) were calculated from the first derivative of the 350/330 nm fluorescence emission ratio. All samples in the sodium acetate buffer systems showed immediate visible precipitation upon addition of LceB and were not included in the nanoDSF experiment.

### Protein crystallization

Crystallization of purified LceB was screened for utilizing commercially available screens (NeXtal) and a Crystal Gryphon robot (Art Robbins Instruments). Crystals of LceB grew in gel filtration buffer and 0.1 M Citric Acid pH 4.0, 10% MPD in a 1:1 at 15 and 10 mg/mL of protein at 4 °C. These conditions were optimized using the sitting drop vapor diffusion method at 4 °C (moved to 20 °C 5 days later), with crystals generated in varying concentrations of citric acid (0.04 M-0.15 M) pH 4.0 and MPD (8-12%) (mother liquor) at a concentration of 10 mg/mL. As the optimized crystals were of insufficient size, they were micro-seeded using the SeedBead kit (Hampton Research). Crystals were obtained in gel filtration buffer 50 mM Tris 750 mM NaCl pH 7.5 at 20 °C from 0.1 M citric acid pH 4.0, 10% MPD at a concentration of 7 mg/mL and used for subsequent data collection and refinement.

### Data Collection and Refinement

The dataset for LceB was collected at the Canadian Light Source beamline CMCF-BM (08B1). Protein crystals were cryo-protected stepwise in 15%, 25% and finally 30% MPD before flash freezing directly in liquid nitrogen. Data was processed using XDS^62^ and CCP4^63^. Initial phases were obtained with Phenix^64^ by molecular replacement utilizing a model of LceB predicted by AlphaFold^65^. The protein was built in Coot^66^ and refined with Phenix^64^ Refmac5^67^ and TLS^68^. UCSF Chimera^69^ and GraphPad Prism version 9.4.1 were utilized for molecular graphics.

### Size exclusion chromatography coupled multiple angle light scattering (SEC-MALS)

Purified LceB was diluted to a concentration of 15 mg/mL in 50 mM TRIS pH 7.5, 750 mM NaCl and spun in a 0.1 μM filter to remove aggregates before SEC-MALS analysis. SEC-MALS data were collected on a DAWN HELEOS II detector (Wyatt Technology) coupled to an AKTA Pure (Cytiva) with an in-line UV cell (Cytiva). For this experiment a Superdex 200 (10/300) increase column (Cytiva) was used. LceB was injected after the system was equilibrated in gel filtration buffer and detectors aligned and normalized with a 5 mg/mL BSA control (Sigma). All experiments were performed at 25 °C. Analysis of the data was completed using ASTRA analysis software (Wyatt Technologies).

### Size exclusion chromatography of digested LceB

Purified LceB was dialyzed O/N at 4 °C with the addition of 1:200 mg/mg LceB:thrombin (Sigma) to remove the N-terminal 6His-tag. After digestion, LceB was first centrifuged to remove precipitation and the soluble fraction was re-purified over an NTA-agarose affinity column to remove undigested material. The flow-through of unbound digested-LceB was concentrated and then analyzed by SEC using an SD75 increase 10/300 column with an AKTAgo (Cytiva) in the optimized buffer (50 mM Tris pH 7.5 750 mM NaCl). Both undigested and digested LceB were analyzed at 5 mg/mL and 1 mg/mL.

### Mass Photometry of LceB

Mass photometry experiments were performed on a Refeyn One Mass Photometer machine (Refeyn Ltd). The protein sample was dialyzed in PBS (pH 7.4). The standard mass photometry landing assay measurements were performed in silicone gaskets on 1.5H high-precision coverslip glass (24×50 mm, Thorlabs). Coverslips were rinsed in the following order: Milli-Q water, isopropanol, and then Milli-Q water. 2 µl of protein sample was added to silicone gaskets containing 15 µl of PBS (pH 7.4) where the final concentration of protein sample was 5 nM. All images were acquired for 1 min at 331 Hz. The measurements were then analyzed and visualized using the Discover MP software (Refeyn Ltd). Measurements were taken in triplicate.

## DATA AVAILABILITY

The X-ray structure and diffraction data reported in this paper have been deposited in the Protein Data Bank under the accession code 8SXQ.

## SUPPORTING INFORMATION

This article contains supporting information.

## ACKNOWLEDGEMENTS

We would like to thank the laboratory of Jörg Stetefeld and Markus Meier at the University of Manitoba for instrument access and assistance with SEC-MALS. Additionally, we thank beamline CMCF-BM at the Canadian Light Source, which is supported by the Canada Foundation for Innovation (CFI), the Natural Sciences and Engineering Research Council (NSERC), the National Research Council (NRC), the Canadian Institutes of Health Research (CIHR), the Government of Saskatchewan.

## FUNDING AND ADDITIONAL INFORMATION

This work was supported by the Natural Sciences and Engineering Research Council of Canada (NSERC) grants RGPIN-2019-05490 to AKB and RGPIN-2018-04968 to G.P., and a Canadian Foundation for Innovation award (CFI) 37841 to G.P. This work was also supported by a University of Manitoba Research Grants Program (URGP) to AKB and GP.

## CONFLICT OF INTEREST

The authors declare that they have no conflicts of interest with the contents of this article

## Notes

### Competing Interest Statement

The authors have declared no competing interest.

