## Supplemental_data for "Structural characterization of the Sel1-like repeat protein LceB from *Legionella pneumophila*"

Running title: LceB is a dynamic Sel-1-like repeat protein

**Key words:**
Effector, LceB, Lpg1356, *Legionella pneumophila*, Type IV secretion system, Sel-1-like Repeat protein, X-ray crystallography

*To whom correspondence should be addressed: G.P.


List of supplementary information:

Figures S1 to S4

Tables S1 to S2

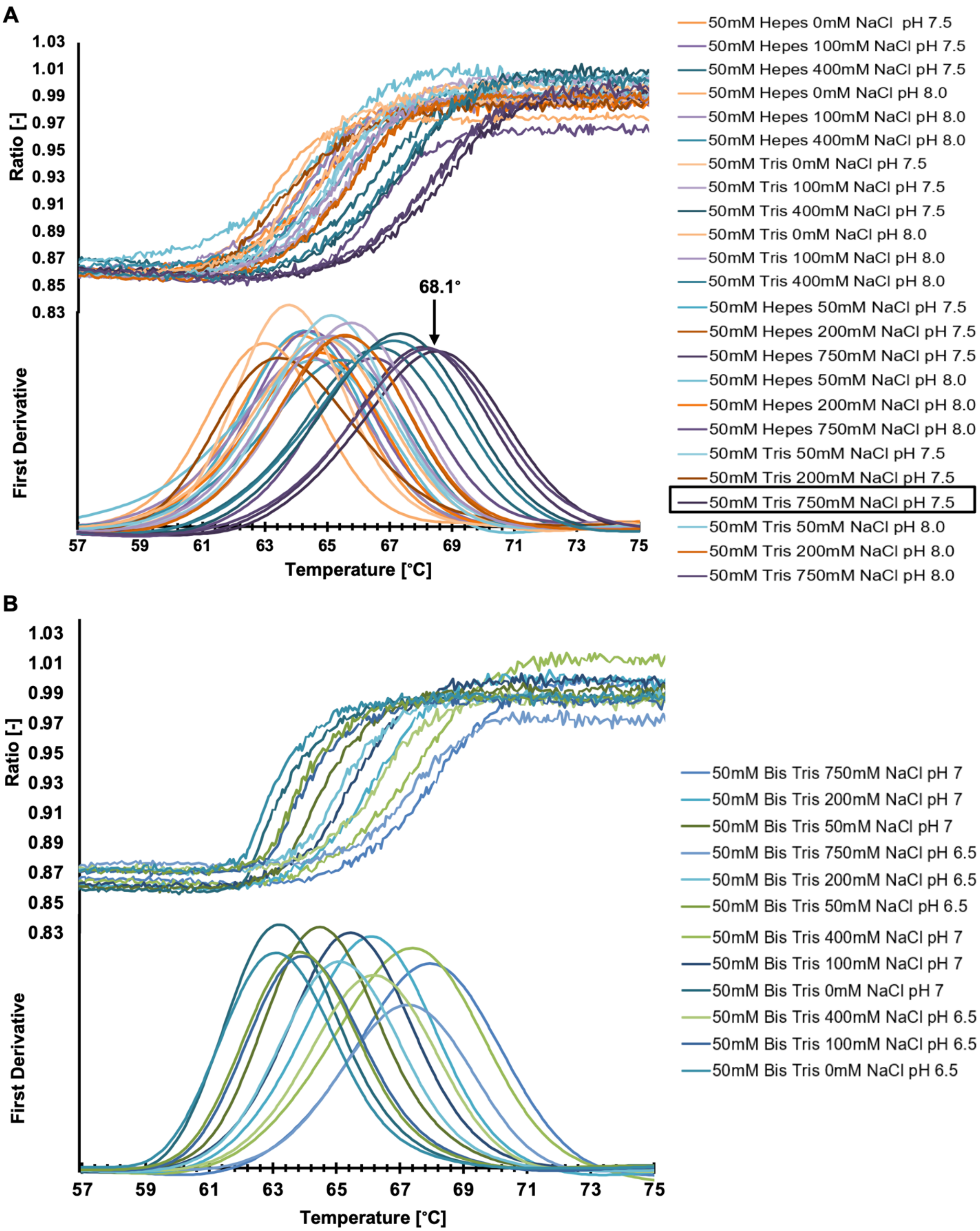

A
Ratio [-]
1.03
1.01
0.99
0.97
0.95
0.93
0.91
0.89
0.87
0.85
0.83
First Derivative
68.1°
57
59
61
63
65
67
69
71
73
75
Temperature [°C]
50mM Hepes 0mM NaCl pH 7.5
50mM Hepes 100mM NaCl pH 7.5
50mM Hepes 400mM NaCl pH 7.5
50mM Hepes 0mM NaCl pH 8.0
50mM Hepes 100mM NaCl pH 8.0
50mM Hepes 400mM NaCl pH 8.0
50mM Tris 0mM NaCl pH 7.5
50mM Tris 100mM NaCl pH 7.5
50mM Tris 400mM NaCl pH 7.5
50mM Tris 0mM NaCl pH 8.0
50mM Tris 100mM NaCl pH 8.0
50mM Tris 400mM NaCl pH 8.0
50mM Hepes 50mM NaCl pH 7.5
50mM Hepes 200mM NaCl pH 7.5
50mM Hepes 750mM NaCl pH 7.5
50mM Hepes 50mM NaCl pH 8.0
50mM Hepes 200mM NaCl pH 8.0
50mM Hepes 750mM NaCl pH 8.0
50mM Tris 50mM NaCl pH 7.5
50mM Tris 200mM NaCl pH 7.5
50mM Tris 750mM NaCl pH 7.5
50mM Tris 50mM NaCl pH 8.0
50mM Tris 200mM NaCl pH 8.0
50mM Tris 750mM NaCl pH 8.0
B
Ratio [-]
1.03
1.01
0.99
0.97
0.95
0.93
0.91
0.89
0.87
0.85
0.83
First Derivative
57
59
61
63
65
67
69
71
73
75
Temperature [°C]
50mM Bis Tris 750mM NaCl pH 7
50mM Bis Tris 200mM NaCl pH 7
50mM Bis Tris 50mM NaCl pH 7
50mM Bis Tris 750mM NaCl pH 6.5
50mM Bis Tris 200mM NaCl pH 6.5
50mM Bis Tris 50mM NaCl pH 6.5
50mM Bis Tris 400mM NaCl pH 7
50mM Bis Tris 100mM NaCl pH 7
50mM Bis Tris 0mM NaCl pH 7
50mM Bis Tris 400mM NaCl pH 6.5
50mM Bis Tris 100mM NaCl pH 6.5
50mM Bis Tris 0mM NaCl pH 6.5

**Figure S1. Thermal denaturation of LceB in different buffer systems**. A) Fluorescence emission ratio for tryptophan at 350/330 nm plotted from 55 ˚C to 75 ˚C of LceB (top panel) and first derivative (bottom panel) after dilution into HEPES and Tris buffer systems. B) Fluorescence emission ratio for tryptophan at 350/330 nm plotted from 55 ˚C to 75 ˚C of LceB (top panel) and first derivative (bottom panel) after dilution into Bis-Tris based buffers. The buffer composition that resulted in the highest protein stability, or melting temperature $T_m$, is boxed and highlighted.

### Table S1: Melting temperatures of LceB in various buffer compositions

| Sample | Tm (C°) | Sample | Tm (C°) |
|---|---|---|---|
| **50mM HEPES pH 7.5 0mM NaCl** | 64.1 | **50mM Tris pH 8.0 0mM NaCl** | 64.8 |
| **50mM HEPES pH 7.5 50mM NaCl** | 64.2 | **50mM Tris pH 8.0 50mM NaCl** | 64.9 |
| **50mM HEPES pH 7.5 100mM NaCl** | 64.5 | **50mM Tris pH 8.0 100mM NaCl** | 65.1 |
| **50mM HEPES pH 7.5 200mM NaCl** | 65.5 | **50mM Tris pH 8.0 200mM NaCl** | 65.5 |
| **50mM HEPES pH 7.5 400mM NaCl** | 66.6 | **50mM Tris pH 8.0 400mM NaCl** | 67.0 |
| **50mM HEPES pH 7.5 750mM NaCl** | 68.0 | **50mM Tris pH 8.0 750mM NaCl** | 68.1 |
| **50mM HEPES pH 8.0 0mM NaCl** | 63.0 | **50mM bis tris 750mM NaCl pH 7** | 67.9 |
| **50mM HEPES pH 8.0 50mM NaCl** | 64.4 | **50mM bis tris 400mM NaCl pH 7** | 67.3 |
| **50mM HEPES pH 8.0 100mM NaCl** | 64.2 | **50mM bis tris 200mM NaCl pH 7** | 66.0 |
| **50mM HEPES pH 8.0 200mM NaCl** | 64.6 | **50mM bis tris 100mM NaCl pH 7** | 65.4 |
| **50mM HEPES pH 8.0 400mM NaCl** | 65.3 | **50mM bis tris 50mM NaCl pH 7** | 64.5 |
| **50mM HEPES pH 8.0 750mM NaCl** | 66.5 | **50mM bis tris 0mM NaCl pH 7** | 63.4 |
| **50mM Tris pH 7.5 0mM NaCl** | 63.8 | **50mM bis tris 750mM NaCl pH 6.5** | 67.2 |
| **50mM Tris pH 7.5 50mM NaCl** | 65.1 | **50mM bis tris 400mM NaCl pH 6.5** | 66.1 |
| **50mM Tris pH 7.5 100mM NaCl** | 65.7 | **50mM bis tris 200mM NaCl pH 6.5** | 65.1 |
| **50mM Tris pH 7.5 200mM NaCl** | 63.6 | **50mM bis tris 100mM NaCl pH 6.5** | 64.1 |
| **50mM Tris pH 7.5 400mM NaCl** | 67.2 | **50mM bis tris 50mM NaCl pH 6.5** | 64.0 |
| **50mM Tris pH 7.5 750mM NaCl** | 68.4 | **50mM bis tris 0mM NaCl pH 6.5** | 63.2 |

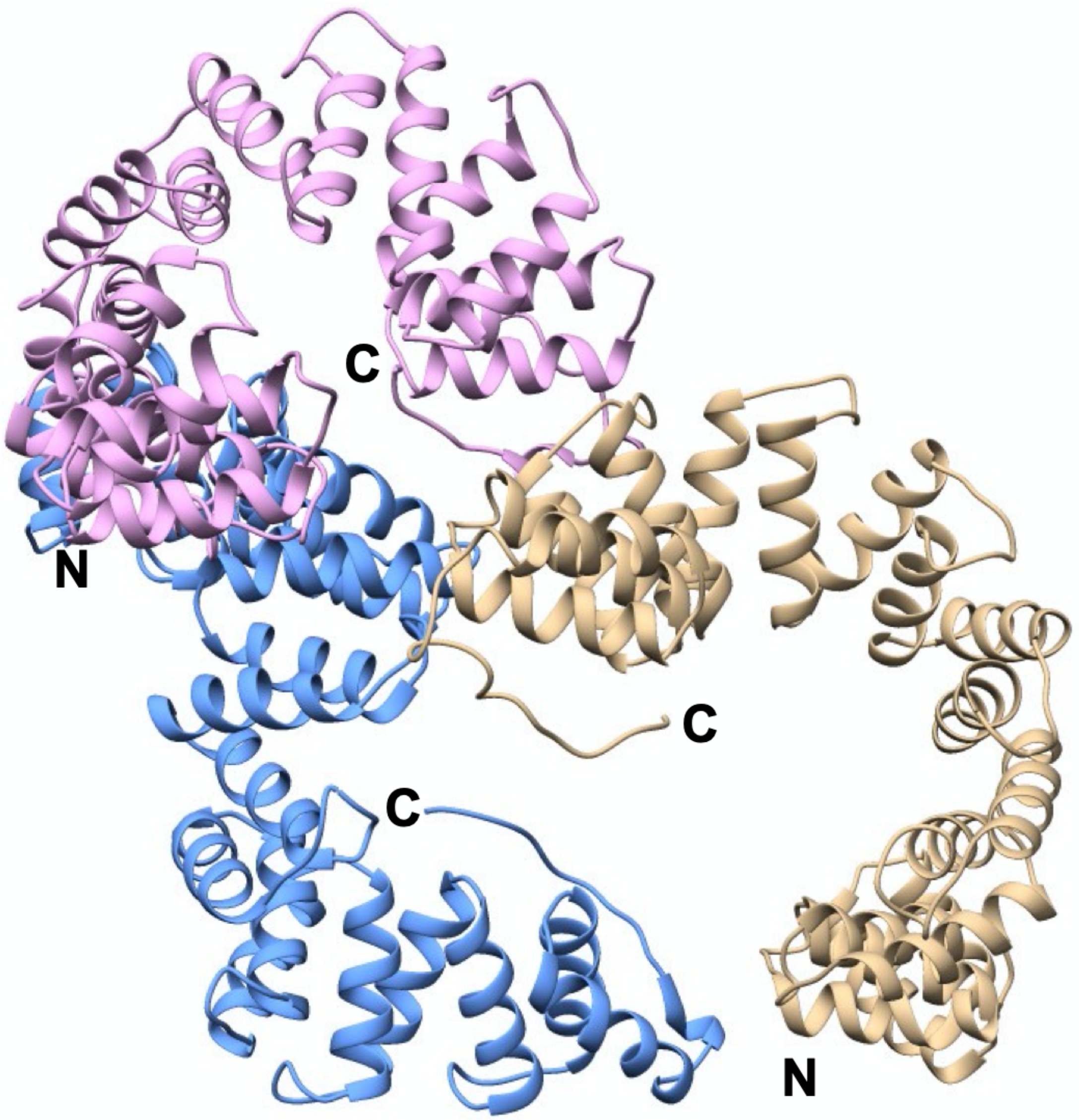


**Figure S2. Configuration of the LceB crystal structure asymmetric unit.** The three LceB chains in the asymmetric unit are depicted in a cartoon ribbon diagram. Chain A (cornflower blue), Chain B (tan), and Chain C (plum). Graphics were drawn using UCSF Chimera (http://www.cgl.ucsf.edu/chimera). N and C termini are indicated.

**Table S2: Analysis of interaction interfaces of the LceB asymmetric unit**

| | Number of residues A | Number of residues C | Number of residues B | Average interface area Å |
|---|---|---|---|---|
| **A and B** | 17 | | 13 | 542.0 |
| **A and C** | 27 | 23 | | 644.3 |
| **B and C** | | 13 | 18 | 578.6 |
| **B and C** | | 13 | 18 | 565.0 |

Interface analysis of asymmetric unit chains A, B, and C by PDBePISA (https://www.ebi.ac.uk/pdbe/pisa/)

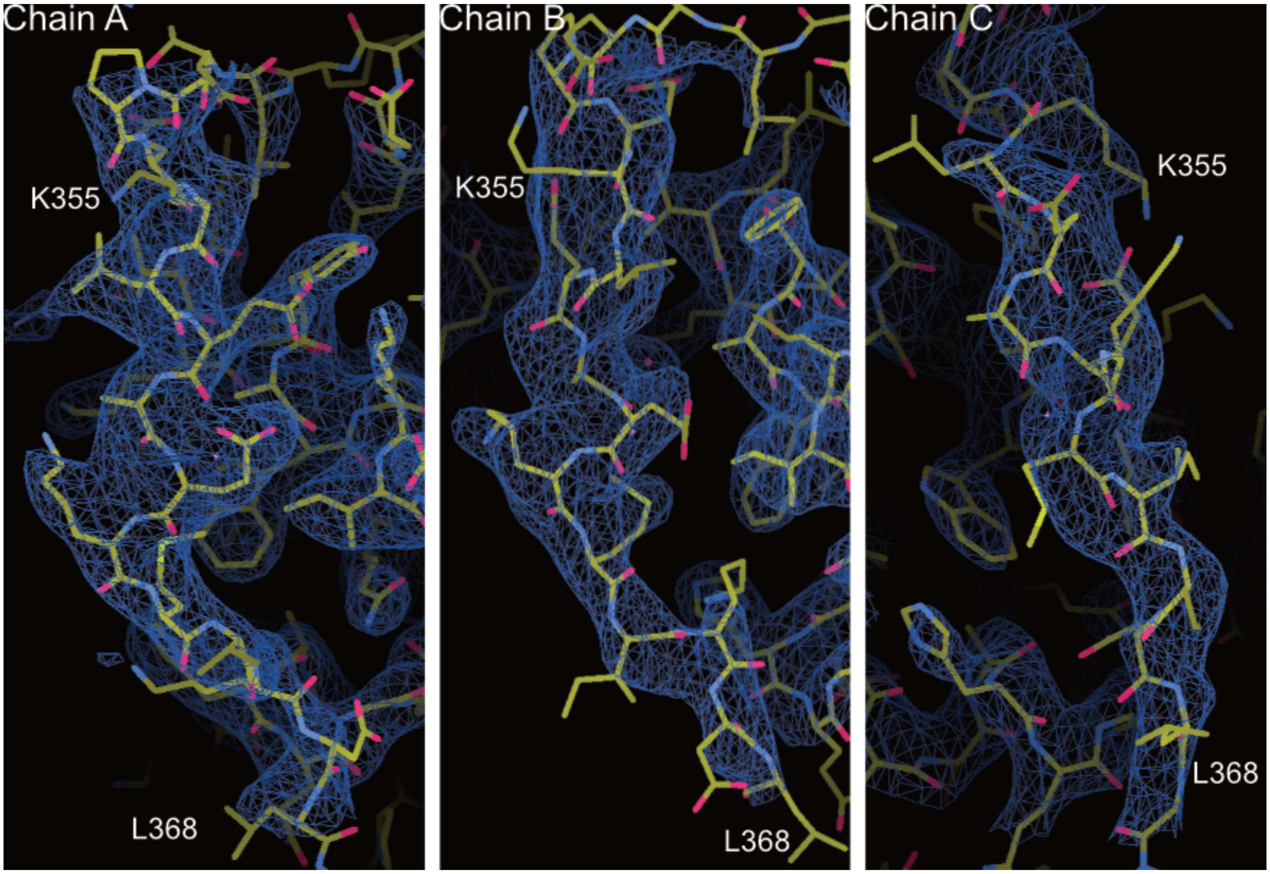


**Figure S3. Electron density maps of the C-terminal helix of LceB.** Electron density maps (*2mFo-DFc*) are shown for restudies K355 to L368 of each chain. Maps are contoured at 1.25 rmsd and drawn using Coot.

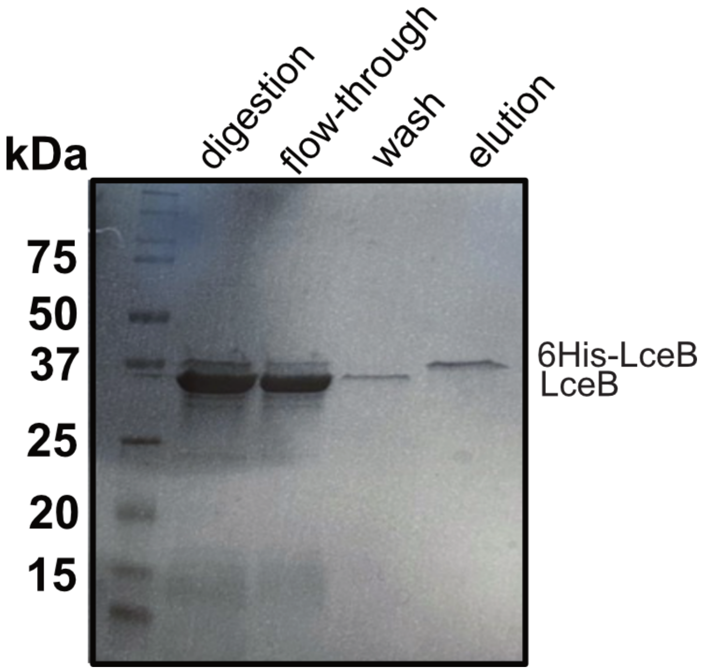


**Figure S4. Proteolytic digestion of 6His-LceB.** Coomassie stained SDS-PAGE gel showing the removal of the N-terminal 6His-tag from LceB. LceB was digested O/N with thrombin at 4˚C and re-purified using a nickel NTA-resin. Lanes are labeled by digestion, flow-through (material that did not bind the NTA-resin), wash (low imidazole buffer wash of resin), and elution (undigested material that still bound to the NTA-resin).
